# p16^INK4A^ expression induces paracrine senescence via small extracellular vesicles

**DOI:** 10.64898/2025.12.09.689258

**Authors:** Marta Menéndez-García, Antonio Merino-Navarro, Ana O’Loghlen

**Affiliations:** Epigenetics & Cellular Senescence Group, Spanish National Research Council (CSIC), Biological Research Centre (CIB), Madrid, Spain

## Abstract

Senescent cells are characterized by the expression of the cell cycle inhibitor and biomarker of aging, p16^INK4A^, and the capacity to modify the microenvironment through the senescence-associated secretory phenotype (SASP). Senescent cells accumulate in physiological and pathological conditions, including aging. In spite of this, fibroblasts ectopically expressing p16^INK4A^ do not release a SASP nor communicate with the microenvironment. Here, we find that human primary fibroblasts expressing p16^INK4A^ release more small extracellular vesicles (sEV) as part of the SASP than proliferating cells. In addition, we show that sEV isolated from p16^INK4A^ cells are able to mediate paracrine senescence by inducing a growth arrest and DNA damage response in proliferating cells albeit not stimulating the expression of IL-8. Furthermore, we show the transmission of paracrine senescence via sEV is conserved in two cellular models of ageing: expression of progerin, mimicking an accelerated form of ageing, and inducing telomere shortening using a dominant negative mutant. Importantly, sEV isolated from fibroblasts derived from old donors also induce paracrine senescence in fibroblasts derived from young donors. In conclusion, our data indicate that sEV released by senescent and aging cells are an important mechanism of intercellular communication and could potentially explain tissue dysfunction in aging.

## INTRODUCTION, RESULTS AND DISCUSSION

Cellular senescence is a process characterized by a stable cell cycle arrest due to upregulation of cyclin kinase inhibitors (Lee & Schmitt, 2019). Senescent cells release inflammatory components (soluble and growth factors, matrix metalloproteinases) collectively named senescence-associated secretory phenotype (SASP) (Coppe et al., 2011). The SASP has an important role in several biological and pathological processes such as wound healing, tissue plasticity, developmental senescence and aging. However, other SASP components such as extracellular vesicles (EV), metabolites, ions and other means of intercellular senescence communication altogether contribute to the tissue homeostasis (Fafian-Labora & O’Loghlen, 2020). EV are membrane particles released by every cell type that contain proteins, nucleic acids and lipids and can be classified by biogenesis or size (e.g. small extracellular vesicles, sEV) (Thery et al., 2018). EV can act as functional secondary messengers reflecting the status of the cell releasing them. Senescent cells release sEV that carry specific DNA, proteins and miR that alter the functionality of the receiving cells (Borghesan et al., 2019; Mensa et al., 2020; Takahashi et al., 2017; Terlecki-Zaniewicz et al., 2018).

*CDKN2A* expression is an important mechanism in the establishment of cellular senescence. It encodes the tumor suppressor and cell cycle inhibitor p16^INK4A^, a major effector of senescence. Thus, p16^INK4A^ has been proposed as a biomarker of aging and senescence both *in vitro* and *in vivo* (He & Sharpless, 2017). Activation of p16^INK4A^ expression induces senescence mimicking different physiological and pathological conditions (Coppe et al., 2011). Thus, the presence of p16^INK4A^ positive cells is frequently observed in most senescent cells, during aging, in progeria syndromes and in benign tumor lesions, e.g. skin nevi, ensuring a stable growth arrest. Importantly, clearance of p16^INK4A^ positive cells prolongs lifespan in mice (Baker et al., 2016; Baker et al., 2011) and attenuates the development of many age-related diseases (He & Sharpless, 2017; Lee & Schmitt, 2019).

Interestingly, Campisi’s group reported that ectopic expression of p16^INK4A^ triggers senescence mimicking aging, without inducing of a SASP (Coppe et al., 2011). As sEV have been recently described to be part of the SASP, we wanted to determine the contribution of sEV isolated from p16^+^ cells taking advantage of human primary fibroblasts (IMR90) ectopically expressing a *CDKN2A* (p16) construct. As a positive control we used IMR90 expressing an inducible 4-hydroxytamoxifen (4OHT) form of oncogenic H-RAS^G12V^ (RAS) or the empty vector (C) (Borghesan et al., 2019; Rapisarda et al., 2017). Thus, we isolated sEV from C, RAS and p16 expressing cells following the differential ultracentrifugation methodology as previously (Borghesan et al., 2019; J. Fafian-Labora et al., 2019) and following the MISEV 2018 guidelines (Thery et al., 2018). The number of particles isolated was analyzed by performing nanoparticle tracking analysis (NTA). Surprisingly, p16^+^ IMR90 released more sEV, which we named evSASP, than C cells (**Fig. 1A**) (Fafian-Labora & O’Loghlen, 2020). As previously published, RAS cells released more sEV than non-senescent cells (Borghesan et al., 2019). In contrast, when analyzing the expression levels of several SASP factors by qPCR, we found that p16^+^ cells did not express what we call a soluble SASP (sSASP), in comparison with RAS-expressing cells (**Fig. 1B**) (Coppe et al., 2011). Importantly, similar findings were reported in fibroblasts derived from the blind mole rat, *Spalax*, undergoing replicative and etoposide-induced senescence where no IL-1α or IL-6 expression could be observed (Odeh, Dronina, Domankevich, Shams, & Manov, 2020). Further characterization of p16^+^ cells confirmed high levels of expression of *CDKN2A* by qPCR (**Fig.1C**), as well as an increase in the mRNA expression levels of the cell cycle inhibitors, *CDKN1A* and *TP53*, in RAS-expressing cells but not in p16^+^ cells (**Fig. 1D**)(Coppe et al., 2011). sEV isolated from p16^+^ cells by NTA analyses show no changes in size distribution between the different groups (**Fig.1E**) and display EV markers as ALIX, TSG1010 and CD9 by immunoblotting (**Fig.1F**). The increase in the release of sEV from p16^+^ cells was further confirmed by capturing sEV to synthetic beads coated with an antibody for a EV biomarker (CD63). Once captured, sEV can be detected by a PE-conjugated secondary antibody for an alternative EV biomarker (CD81) by flow cytometry. As shown in **Fig.1G**, p16^+^ IMR90 release more sEV than their proliferating counterparts. These data are in accordance with several studies showing an increase in the release of sEV from cells undergoing senescence independent of the trigger (Borghesan et al., 2019; J. Fafian-Labora et al., 2019; Jeon et al., 2019; Takahashi et al., 2017; Terlecki-Zaniewicz et al., 2018).

**Figure 1.**
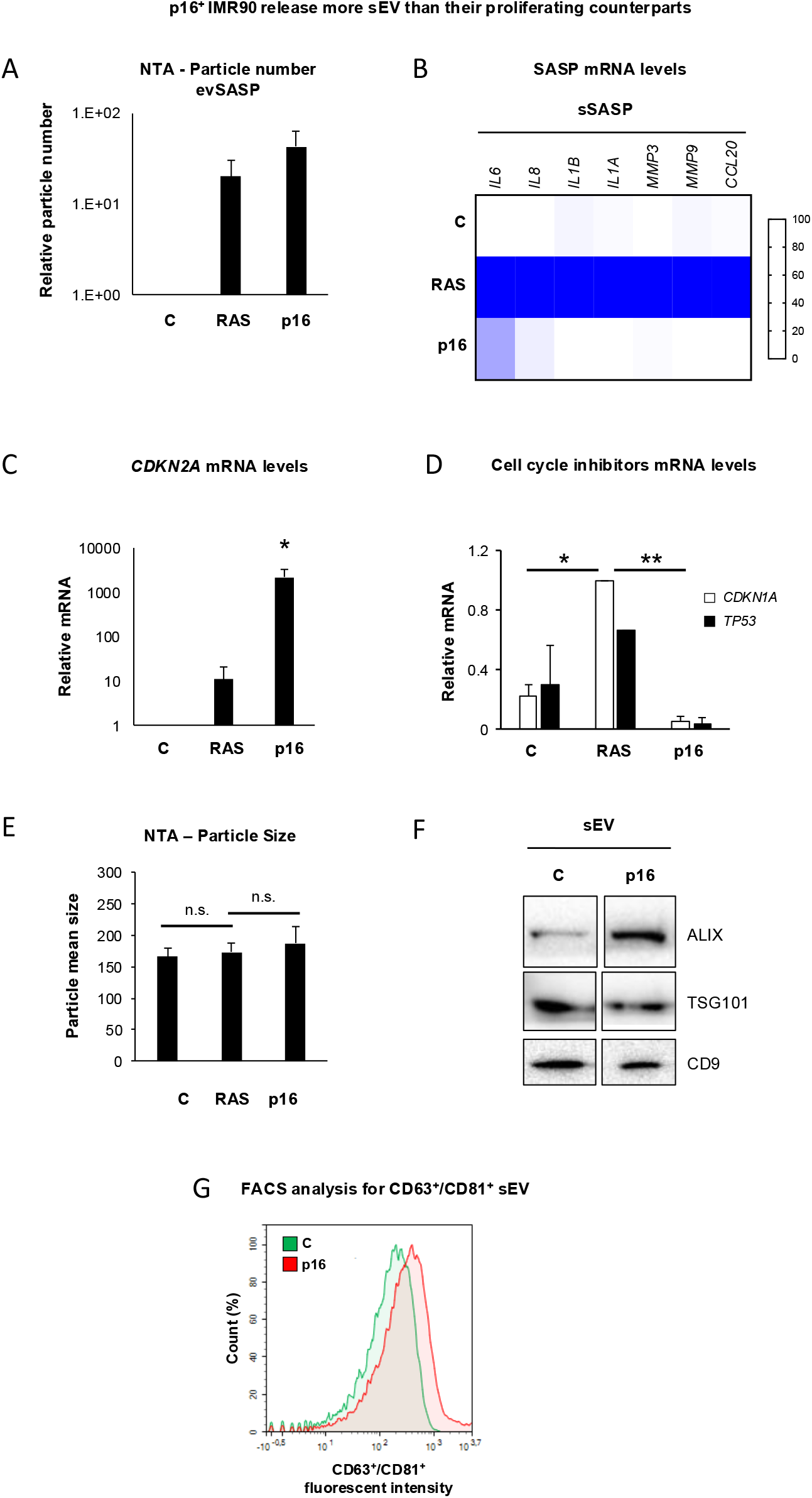
p16^INK4A^-expressing IMR90 release more sEV than proliferating IMR90 cells. **(A)** NTA analysis measuring the number of sEV released by IMR90 infected with control vector (C), inducible vector encoding ER:H-RAS^G12V^ (RAS) and ectopic expression of p16^INK4A^ (p16). Data represent the mean ± SEM of three independent experiments. **(B)** Heatmap showing the relative mRNA levels related to SASP genes in C, RAS and p16 cells by qPCR. The data show the mean from three independent experiments. **(C)** Quantification of *CDKN2A* mRNA levels by qPCR show an increase when p16 is ectopically expressed. RAS cells were used as a positive control. **(D)** *CDKN1A* and *TP53* mRNA levels extracted from C, RAS and p16 are shown. **(C, D)** Data show the mean ± SEM of 3 independent experiments. Student’s t-test was performed. **(E)** NTA analyses measuring particle size in sEV isolated from C, RAS and p16 cells. Data represent the mean ± SEM of 3 experiments. Student’s t-test. **(F)** Representative immunoblot for sEV markers (ALIX, TSG101 and CD9) in sEV isolated from C and p16 IMR90. **(G)** FACS histogram of CD63^+^/CD81^+^ sEV released from C and p16 IMR90 and captured to latex beads. Representative experiment is shown.

We and others have identified that evSASP can mediate paracrine senescence in different contexts (Borghesan et al., 2019; Jeon et al., 2019; Mensa et al., 2020). To determine if sEV isolated from p16^+^ IMR90 were involved in mediating paracrine senescence, we treated proliferative IMR90 cells with sEV isolated from C, RAS and p16 IMR90 cells. First, we determined whether p16^+^ sEV altered the proliferative capacity of IMR90 recipient cells by quantifying the percentage of cells staining positive for Ki-67 by immunofluorescence (**Fig.2A**). Next, we show that sEV from p16^+^ cells induced an increase in the percentage of cells staining positive for p16^INK4A^ (**Fig.2B**). The cell cycle arrest was concomitant with an increase in the number of cells staining positive for γ-pH2AX, a maker of DNA damage (**Fig.2C**). In contrast, we find the percentage of cells staining positive for IL-8 is low in a high number of recipient cells treated with sEV from p16^+^ cells, in contrast with cells treated with sEV from iRAS cells (**Fig.2D**). Altogether, our data suggest that p16^+^ cells release evSASP but not sSASP. In addition, our data show the evSASP released by p16^+^ cells have the ability to induce paracrine senescence, although in contrast sEV from iRAS cells they do not induce the expression of IL-8 in particular. Next, we generated an alternative cellular model mimicking aging by ectopically expressing either the dominant-negative allele of the telomeric repeat binding factor 2 (TRF2^ΔBΔM^) or an inducible construct expressing progerin (PG) in IMR90 cells. We analyzed whether sEV isolated from these aging models induced paracrine evSASP in IMR90 by measuring the percentage of cells incorporating BrdU (**Fig.2E**). Both cellular models were capable of inducing paracrine senescence mediated by sEV. As p16^+^ cells are enriched during aging, we next took advantage of a human fibroblast culture isolated from a 1 year-old donor GM05399 and treated it with sEV isolated from a 67 year-old human fibroblast culture (AG16086). By measuring BrdU we could observe that sEV from AG16086 induced a growth arrest in GM05399 confirming transfer of paracrine senescence during aging (**Fig.2F**). Finally, we isolated sEV from four different fibroblasts derived from old donors (∼70 years) and treated individually four fibroblasts from young donors (∼2 years) and measured β-gal activity by IF. Our data show an increase in β-gal activity in young cells when treated with evSASP from old donors (**Fig.2G**). Altogether, using different cellular models of aging and cells derived from young donors we show that evSASP mediates paracrine senescence, even when the trigger induces senescence without a sSASP.

**Figure 2.**
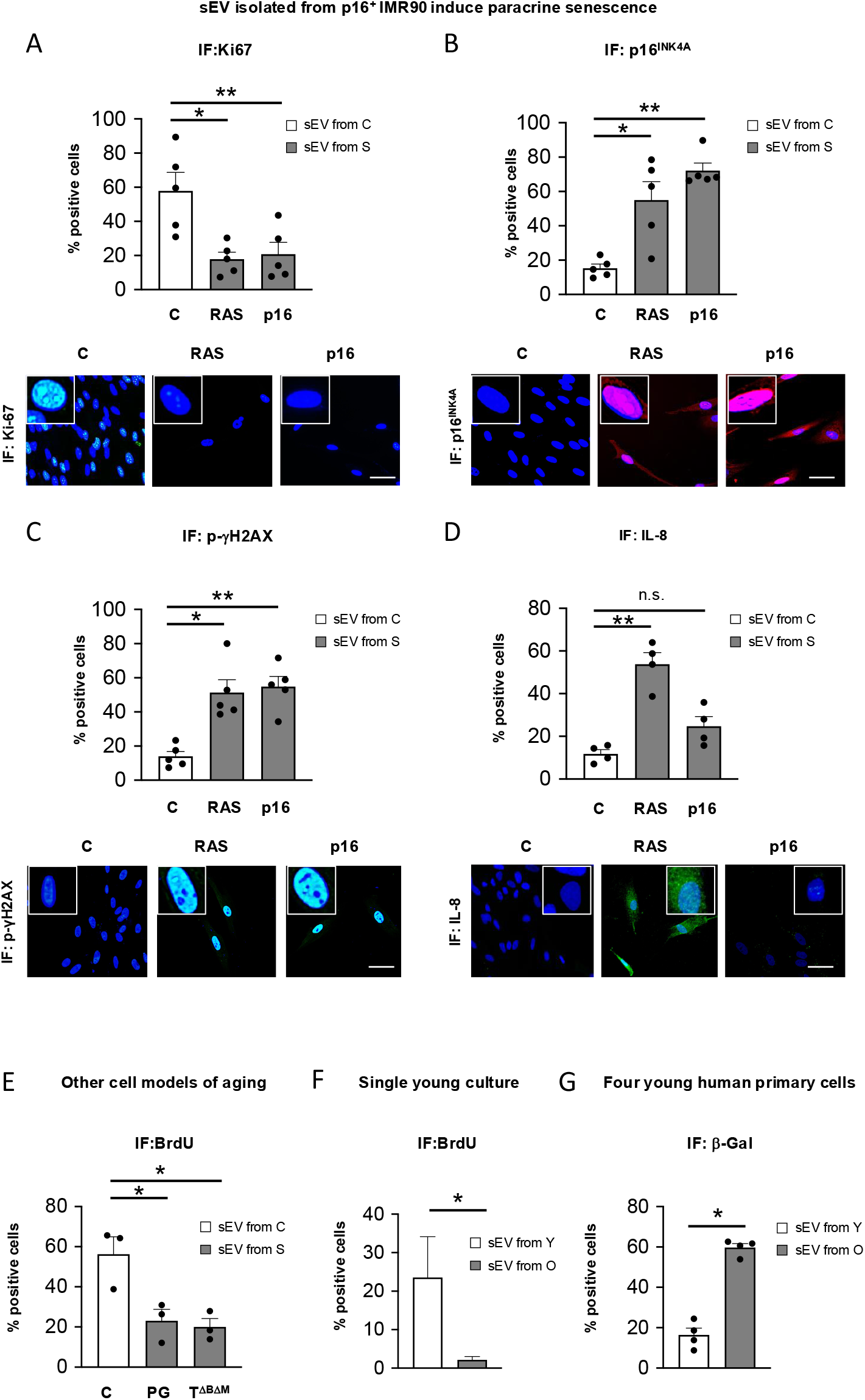
sEV isolated from p16^INK4A^ -IMR90 cells induce paracrine senescence in proliferating IMR90. **(A-D)** Immunofluorescence quantification (top panels) and representative images (bottom panels) for different biomarkers of senescence: **(A)** Ki67, **(B)** p16^INK4A^, **(C)** p-γH2AX, **(D)** IL-8 in IMR90 cells treated with sEV from C or sEV from senescent (S) cells expressing either RAS or p16 for 72h. Scale bar, 50 μM. Data represent the mean ± SEM of 4-5 independent experiments. One-way ANOVA analysis. **(E)** Quantification of the percentage of cells incorporating BrdU in IMR90 treated with sEV from C or from senescent (S) IMR90 expressing progerin (PG) or TRF2^ΔDΔM^ (T^ΔDΔM^) constructs. Data represent the mean ± SEM of 3 independent experiments. One-way ANOVA analysis. **(F)** Young fibroblast culture GM05399 was treated with sEV isolated from the old culture AG16086 for 72h. BrdU incorporation was measured. Data show the mean ± SEM of 3 experiments. Student’s t-test analysis. **(G)** Four young human primary fibroblasts were treated for 72h with sEV isolated from old primary fibroblasts and β-Gal activity was measured by IF. Data represent the mean ± SEM of 4 individual young cells. t-test analysis was performed.

A few decades ago, it was thought that tissues aged due to their lack of proliferative and regenerative capacity (He & Sharpless, 2017) and that p16^+^ cells did not communicate with the microenvironment (Coppe et al., 2011). At present, it is acknowledged that the elimination of senescent cells by the recruitment of the immune system through sSASP is key, mechanism that is compromised during ageing. One hypothesis based on our data could be that p16^+^ senescent cells that are enriched during ageing have a weak or non-existent initial acute sSASP activation. Thus, they induce paracrine senescence via evSASP to reinforce the recall to the immune system. In addition, the induction of paracrine senescence through evSASP enhances the lack of proliferative capacity of the tissue by damaging healthy cells which arrest but do not induce sSASP. These findings highlight the importance of understanding different means of non-cell autonomous intercellular communication and the influence it has on the microenvironment in senescence and aging.

## ACKNOWLEDGEMENTS

AO’s lab is supported by BBSRC (BB/P000223/1), Barts Charity Grants (MGU0497, G002158), Project I+D+i PID2021-125656OB-I00 and CNS2022-135134, financed by MICIU/AEI/10.13039/501100011033/”ERDF/EU, and Project SenesceX-CM P2022/ BMD-7393. MMG received a PhD fellowship from Comunidad de Madrid (PIPF-2023/SAL-GL-29915).

## DATA AVAILABILITY STATEMENT

The data that support the findings of this study are available from the corresponding author, AOL, upon reasonable request.

## EXPERIMENTAL PROCEDURES

### Cell Culture

IMR90 and HEK293T cells were obtained from the American Type Culture Collection (ATCC). Cells were cultured in Dulbecco’s modified Eagle’s medium (DMEM) supplemented with 10 % (v/v) fetal bovine serum (FBS) and 1 % (v/v) antibiotic-antimycotic (A/A) solution (all from Sigma-Aldrich). To collect small extracellular vesicles (sEV), the cells were maintained in DMEM supplemented with 0.5 % (v/v) sEV-depleted FBS and 1 % (v/v) A/A for 72 h. FBS was depleted of sEV by overnight ultracentrifugation at 100,000 xg at 4C. Young and old-derived human primary fibroblasts were obtained from the Coriell Cell Repository with the following codes: Young: GM05399 (1 year old; Y1), GM00969 (2 years old; Y2), GM05565 (3 years old; Y3), GM05758 (1 year old; Y4); Old: AG16086 (67 years old; O1), AG06240 (80 years old; O2), AG13152 (80 years old; O3), AG13222 (81 years old; O4).

### Retroviral infections

For retroviral production, HEK293T cells were cotransfected with vsv-g envelope plasmid and Gag-Pol helper vector using polyethylenimine linear (Sigma-Aldrich). The viral supernatant was collected from the HEK293T cells 2 days after transfection. It was filtered 0.45-μm syringe filter (Starlab) to eliminate cells. The filtered viral supernatant was supplemented with hexadimethrine bromide (4 μg/ml; polybrene) (Sigma-Aldrich) and used to infect IMR90 cells overnight, with two subsequent collections of viral supernatant and treatments for 3 hours each. After three rounds of infection, the medium was changed to fresh DMEM supplemented with 10% (v/v) FBS and 1 % (v/v) A/A. After two days, IMR90 cells were selected with DMEM supplemented with 10% (v/v) FBS, 1 % (v/v) A/A and 0.5 μg/ml puromycin or 300 μg/ml neomycin (Invitrogen) for one week before starting the experiments.

### Isolation of sEV and treatment of recipient cells

Conditioned medium (CM) from 10^6^ IMR90 cultured in 0.5% EV-depleted FBS media for 72h and plated in a 10cm dish was collected. CM was centrifuged at low speed to eliminate dead cells, filtered (0.22µm) and centrifuged at 10,000xg for 1h. The pellet was washed with 15ml PBS at 100,00xg for 1h 20min. It was resuspended in 100µl of media and recipient IMR90 were incubated with sEV in media supplemented with 10% EV-depleted FBS for 72 h. sEV obtained from 100mm plates were used to treat 12 wells of a 96 well-plate (Corning). sEV obtained from all 4 human old primary fibroblast donors were mixed and used to treat individually 4 young recipient cells maintaining the previous proportion of sEV isolated from one 10cm dish used to treated 12 wells of a 96-well plate. For Western Blot analysis, sEV were resuspended in protein lysis buffer. A Sorvall 100SE Ultra Centrifuge, with a Beckmann Fixed Angle T865 rotor was used.

### Nanoparticle tracking analysis

The NanoSight LM10 (Malvern Instruments, Malvern, United Kingdom) was used to determine number and size of particles. The machine was calibrated using Silica Microspheres beads (Polyscience, Illinois, USA) before measurements were taken. The preparation of sEV were diluted in PBS (Sigma-Aldrich, MO, USA) until obtaining a concentration of particle number between 10^8^ –10^9^ particles. Three measurements of 60 s were taken per each sample and the mean value was used to determine particle number. The NTA 3.0 software (Malvern Instruments, Malvern, United Kingdom) was used to generate the average displacement of each particle per unit second from the movement of each particle in the field of views.

### FACS beads capture

For the determination of CD63/CD81 positive sEV by FACs, aldehyde/sulphate latex beads (ThermoFisher) were coated with an anti-CD63 antibody overnight. Afterwards, the antibody-coated beads were washed three times with PBS. Then, they were incubated the different conditioned medium overnight at 4C in a rotation wheel. After extensive washing with PBS, the sEV-beads complexes were incubated with anti-CD81-PE conjugated antibody for 1h at RT in a rotation wheel, washing three times with PBS and acquired using NovoCyte Flow Cytometer (Acea, Biosciences). Details of antibodies used: CD81-PE (1/200; Life Technologies Cat# A15781), CD63 (1/100; BD Cat# 556019).

### Immunoblot

sEV were lysed using the following lysis buffer [(Tris-HCl 20 mM pH 7.6; DTT 1 mM; EDTA 1 mM; PMSF 1 mM; benzamidine 1 mM; sodium molybdate 2 mM; β-sodium glycerophosphate 2 mM; sodium orthovanadate 0.2 mM; KCl 120 mM; 1 mg/ml (each) leupeptin, pepstatin A and antipain; NonidetTM P-40 0.5% (v/v); Triton X-100 0,1% (v/v)] (all from Sigma-Aldrich).

Lysates were diluted with 4X Laemmli Sample Buffer (Bio-Rad, UK) and an equal number of particles (3×10^9^) were separated in SDS-PAGE gels at 80V. Then, they were transferred to a PVDF 0.45 mm pore size membrane using wet transfect at 350 mA for 2 h at 4C. The membranes were blocked using 5 % (m/v) BSA in 0.1 % (v/v) PBS-Tween for 1 h. After, they were incubated with different antibodies as detailed below. Protein bands were detected using a SuperSignal West Pico PLUS Chemiluminescent Substrate and the ChemiDoc XRS+ System (Bio-Rad). Primary antibodies used: TSG101 (1/1000; Abcam Cat# ab30871), ALIX (1/1000; Abcam Cat# ab88743), CD9 (1/1000; Abcam Cat# ab223052).

### qPCR

Total RNA was extracted using Trizol Reagent (Invitrogen). cDNA was synthesized using the High Capacity cDNA Reverse transcription kit (Invitrogen). qPCR was performed using SYBR Green PCR Master Mix (Applied Biosystems) on a 7500 Fast System RealTime PCR cycler (Applied Biosystems). Relative gene expression was normalized to *RPS14*. The primers used in this study are:

*CDKN2A*: For CGGTCGGAGGCCGATCCAG; Rev GCGCCGTGGAGCAGCAGCAGCT

*CDKN1A*: For CCTGTCACTGTCTTGTACCCT; Rev GCGTTTGGAGTGGTAGAAATC

*TP53*: For CCGCAGTCAGATCCTAGCG; Rev AATCATCCATTGCTTGGGACG

*IL6*: For CCAGGAGCCCAGCTATGAAC; Rev CCCAGGGAGAAGGCAACTG

*IL8*: For GAGTGGACCACACTGCGCCA; Rev TCCACAACCCTCTGCACCCAGT

*IL1B*: For TGCACGCTCCGGGACTCACA; Rev CATGGAGAACACCACTTGT

*IL1A*: For AGTGCTGCTGAAGGAGATGCCTGA; Rev CCCCTGCCAAGCACACCCAGTA

*MMP3*: For AAGCTCTGAAAGTCTGGGAAGA; Rev TCCCTGTTGTATCCTTTGTCCA

*MMP9*: For AACTTTGACAGCGACAAGAAGT; Rev ATTCACGTCGTCCTTATGCAAG *CCL20*: For GGCGAATCAGAAGCAGCAAGCAAC; Rev ATTGGCCAGCTGCCGTGTGAA

*RPS14*: For CTGCGAGTGCTGTCAGAGG; Rev TCACCGCCCTACACATCAAACT

### Immunofluorescence and high content analysis

IMR90 cells were washed with PBS three times, fixed in 4% (v/v) paraformaldehyde (Sigma-Aldrich) for 15min at room temperature and permeabilized with 0.2 % (v/v) Triton X-100 (Sigma-Aldrich) for 20min at room temperature. Plates were blocked with 1% (m/v) BSA and 0.2 % (m/v) gelatin fish (all from Sigma-Aldrich) and incubated with the primary antibody overnight at 4C followed by the secondary antibody and DAPI. Images were acquired and analyzed using the automated fluorescent microscope INCell Analyzer and Investigator 2200 (GE Healthcare) taking 12 fields per well. Primary antibodies used: p16^INK4A^ (1/500; Cat# ab108349), phospho-γH2AX (1/500; Merck Millipore Cat# 05-636-I), IL-8 (1/500; R&D Systems Cat# MAB208), Ki67 (1/500; Abcam Cat# ab92742,), BrdU 1/500 (Abcam Cat # ab6326). For β-Galactosidase staining by IF, IMR90 were incubated with 33 µM of the β-galactosidase substrate C12FDG (Fluorescein di-β-D-galactopyranoside; Sigma-Aldrich F2756) in medium supplemented with 0.5 % (v/v) FBS-depleted sEV for 8h at 37C. After incubation, cells were washed with PBS and fixed with 4% (v/v) paraformaldehyde for 15 min at room temperature.

### Statistics analysis

The results are expressed as the mean ± SEM unless specified and statistical analysis was performed using a Student’s t test unless specified. A p < 0.05 was considered significant. *p< 0.05; **p < 0.01; *** p < 0.001

## REFERENCES

Acosta, J. C., O’Loghlen, A., Banito, A., Raguz, S., & Gil, J. (2008). Control of senescence by CXCR2 and its ligands. Cell Cycle, 7(19), 2956–2959. doi:10.4161/cc.7.19.6780

Baker, D. J., Childs, B. G., Durik, M., Wijers, M. E., Sieben, C. J., Zhong, J., … van Deursen, J. M. (2016). Naturally occurring p16(Ink4a)-positive cells shorten healthy lifespan. Nature, 530(7589), 184–189. doi:10.1038/nature16932

Baker, D. J., Wijshake, T., Tchkonia, T., LeBrasseur, N. K., Childs, B. G., van de Sluis, B., … van Deursen, J. M. (2011). Clearance of p16Ink4a-positive senescent cells delays ageing-associated disorders. Nature, 479(7372), 232–236. doi:10.1038/nature10600

Basisty, N., Kale, A., Jeon, O. H., Kuehnemann, C., Payne, T., Rao, C., … Schilling, B. (2020). A proteomic atlas of senescence-associated secretomes for aging biomarker development. PLoS Biol, 18(1), e3000599. doi:10.1371/journal.pbio.3000599

Borghesan, M., Fafian-Labora, J., Eleftheriadou, O., Carpintero-Fernandez, P., Paez-Ribes, M., Vizcay-Barrena, G., … O’Loghlen, A. (2019). Small Extracellular Vesicles Are Key Regulators of Non-cell Autonomous Intercellular Communication in Senescence via the Interferon Protein IFITM3. Cell Rep, 27(13), 3956–3971 e3956. doi:10.1016/j.celrep.2019.05.095

Coppe, J. P., Rodier, F., Patil, C. K., Freund, A., Desprez, P. Y., & Campisi, J. (2011). Tumor suppressor and aging biomarker p16(INK4a) induces cellular senescence without the associated inflammatory secretory phenotype. J Biol Chem, 286(42), 36396–36403. doi:10.1074/jbc.M111.257071

Demaria, M. (2018). Gene therapy for p16-overexpressing cells. Aging (Albany NY), 10(4), 518–519. doi:10.18632/aging.101422

Fafian-Labora, J., & O’Loghlen, A. (2020). Classical and non-classical intercellular communication in senescence and ageing. Trends Cell Biol.

Mensa, E., Guescini, M., Giuliani, A., Bacalini, M. G., Ramini, D., Corleone, G., … Olivieri, F. (2020). Small extracellular vesicles deliver miR-21 and miR-217 as pro-senescence effectors to endothelial cells. J Extracell Vesicles, 9(1), 1725285. doi:10.1080/20013078.2020.1725285

O’Loghlen, A. (2018). Role for extracellular vesicles in the tumour microenvironment. Philos Trans R Soc Lond B Biol Sci, 373(1737). doi:10.1098/rstb.2016.0488

Odeh, A., Dronina, M., Domankevich, V., Shams, I., & Manov, I. (2020). Downregulation of the inflammatory network in senescent fibroblasts and aging tissues of the long-lived and cancer-resistant subterranean wild rodent, Spalax. Aging Cell, 19(1), e13045. doi:10.1111/acel.13045

Paez-Ribes, M., Gonzalez-Gualda, E., Doherty, G. J., & Munoz-Espin, D. (2019). Targeting senescent cells in translational medicine. EMBO Mol Med, 11(12), e10234. doi:10.15252/emmm.201810234

Patil, P., Dong, Q., Wang, D., Chang, J., Wiley, C., Demaria, M., … Vo, N. (2019). Systemic clearance of p16(INK4a) -positive senescent cells mitigates age-associated intervertebral disc degeneration. Aging Cell, 18(3), e12927. doi:10.1111/acel.12927

Rapisarda, V., Borghesan, M., Miguela, V., Encheva, V., Snijders, A. P., Lujambio, A., & O’Loghlen, A. (2017). Integrin Beta 3 Regulates Cellular Senescence by Activating the TGF-beta Pathway. Cell Rep, 18(10), 2480–2493. doi:10.1016/j.celrep.2017.02.012

Sessions, G. A., Copp, M. E., Liu, J. Y., Sinkler, M. A., D’Costa, S., & Diekman, B. O. (2019). Controlled induction and targeted elimination of p16(INK4a)-expressing chondrocytes in cartilage explant culture. FASEB J, 33(11), 12364–12373. doi:10.1096/fj.201900815RR

Takahashi, A., Okada, R., Nagao, K., Kawamata, Y., Hanyu, A., Yoshimoto, S., … Hara, E. (2018). Publisher Correction: Exosomes maintain cellular homeostasis by excreting harmful DNA from cells. Nat Commun, 9(1), 4109. doi:10.1038/s41467-018-06613-3

Terlecki-Zaniewicz, L., Lammermann, I., Latreille, J., Bobbili, M. R., Pils, V., Schosserer, M., … Grillari, J. (2018). Small extracellular vesicles and their miRNA cargo are anti-apoptotic members of the senescence-associated secretory phenotype. Aging (Albany NY), 10(5), 1103–1132. doi:10.18632/aging.101452

Thery, C., Witwer, K. W., Aikawa, E., Alcaraz, M. J., Anderson, J. D., Andriantsitohaina, R., … Zuba-Surma, E. K. (2018). Minimal information for studies of extracellular vesicles 2018 (MISEV2018): a position statement of the International Society for Extracellular Vesicles and update of the MISEV2014 guidelines. J Extracell Vesicles, 7(1), 1535750. doi:10.1080/20013078.2018.1535750

Fafian-Labora, & O’Loghlen, A. (2020). Classical and non-classical intercellular communication in senescence and ageing. Trends Cell Biol.

Fafian-Labora, J., Carpintero-Fernandez, P., Jordan, S. J. D., Shikh-Bahaei, T., Abdullah, S. M., Mahenthiran, M., … O’Loghlen, A. (2019). FASN activity is important for the initial stages of the induction of senescence. Cell Death Dis, 10(4), 318. doi:10.1038/s41419-019-1550-0

He, S., & Sharpless, N. E. (2017). Senescence in Health and Disease. Cell, 169(6), 1000–1011. doi:10.1016/j.cell.2017.05.015

Jeon, O. H., Wilson, D. R., Clement, C. C., Rathod, S., Cherry, C., Powell, B., … Elisseeff, J. H. (2019). Senescence cell-associated extracellular vesicles serve as osteoarthritis disease and therapeutic markers. JCI Insight, 4(7). doi:10.1172/jci.insight.125019

Lee, S., & Schmitt, C. A. (2019). The dynamic nature of senescence in cancer. Nat Cell Biol, 21(1), 94–101. doi:10.1038/s41556-018-0249-2

Takahashi, A., Okada, R., Nagao, K., Kawamata, Y., Hanyu, A., Yoshimoto, S., … Hara, E. (2017). Exosomes maintain cellular homeostasis by excreting harmful DNA from cells. Nat Commun, 8, 15287. doi:10.1038/ncomms15287

